# Is evolution in response to extreme events good for population persistence?

**DOI:** 10.1101/2020.04.02.014951

**Authors:** Kelsey Lyberger, Matthew Osmond, Sebastian Schreiber

## Abstract

Climate change is predicted to increase the severity of environmental perturbations, including storms and droughts, which act as strong selective agents. These extreme events are often of finite duration (pulse disturbances). Hence, while evolution during an extreme event may be adaptive, the resulting phenotypic changes may become maladaptive when the event ends. Using individual-based models and analytic approximations that fuse quantitative genetics and demography, we explore how heritability and phenotypic variance affect population size and extinction risk in finite populations under an extreme event of fixed duration. Since more evolution leads to greater maladaptation and slower population recovery following an extreme event, slowing population recovery, greater heritability can increase extinction risk when the extreme event is short, as in random environments with low autocorrelation. Alternatively, when an extreme event is sufficiently long, heritability often helps a population persist, as under a press perturbation. We also find that when events are severe, the buffering effect of phenotypic variance can outweigh the increased load it causes. Our results highlight the importance of the length and severity of a disturbance when assessing the role of evolution on population recovery; the rapid adaptive evolution observed during extreme events may be bad for persistence.

## Introduction

Globally, humans are causing substantial environmental perturbations, and these perturbations are likely to become more severe in the future. In particular, climate change is projected to lead to more extreme weather events, including droughts and major storms (Ummenhofer and Meehl, 2017). With more severe events comes the potential for dramatic demographic and genetic consequences.

In the process of causing mass mortality, extreme events can act as catalysts of evolutionary change. In fact, there are many examples of rapid evolution in response to extreme events (reviewed in Grant et al., 2017). Famously, Bumpus (1899) documented phenotypic differences in house sparrows that survived a strong winter storm. More recently, Donihue et al. (2018) measured lizards before and after a series of hurricanes and found evidence for selection on body size, relative limb length, and toepad size. Another example is a study of the annual plant *Brassica rapa* in response to summer drought, in which, post-drought seeds flowered earlier when planted alongside pre-drought seeds (Franks et al., 2007). Finally, Grant and Grant (2014) not only documented shifts in beak depth of Darwin’s ground finches in response to drought, but also the reversal of that evolution and population recovery in subsequent years. We have many fewer examples like this latter case, where the recovery from an extreme event is recorded. Hence exploring what factors influence recovery patterns is currently best done with theory.

Extreme events such as storms, hurricanes, and droughts are pulse disturbances, defined as a transient or short-term change in the environment (Bender et al., 1984). In the ecological literature, pulse disturbances lie at the crossroads of two other forms of environmental change: press perturbations, a permanent or long-term change in the environment (Ives and Carpenter, 2007; Kéfi et al., 2019; Yodzis, 1988), and fluctuating environments such as a sequence of pulse disturbances of varying durations and intensities (Lande, 1993; Ozgul et al., 2012). Despite their transient nature, pulse disturbances can substantially impact ecological systems from large, long-term transients in ecological dynamics to permanent shifts in ecological states (Fox and Gurevitch, 2000; Hastings et al., 2018; Holling, 1973, 1996; Holt, 2008; Ives and Carpenter, 2007). While the most extreme form of a permanent shift is species extinction, we know of no studies that examine the effects of pulse disturbances of varying length on extinction risk (see, however, Figure 1 in Holt, 2008). In contrast, there have been many studies examining the effect of repeated disturbances on extinction risk (Cuddington and Yodzis, 1999; Mangel and Tier, 1994; Petchey et al., 1997). For example, Wilson and MacArthur (1967) showed that the mean time to extinction plateaus as a function of the initial population size. This plateau, which allows one to define a minimum viable population size, is lost when populations experience repeated large disturbances; there is a continual gradual increase in the mean time to extinction with increasing initial population sizes (Mangel and Tier, 1994). In these studies of repeated disturbances, extinction occurs eventually. Consequently, the focus is on when the extinction event occurs, not on whether it occurs after a particular disturbance. However, understanding short-term exinction risk after a single disturbance is critical for conservation and management.

**Figure 1:**
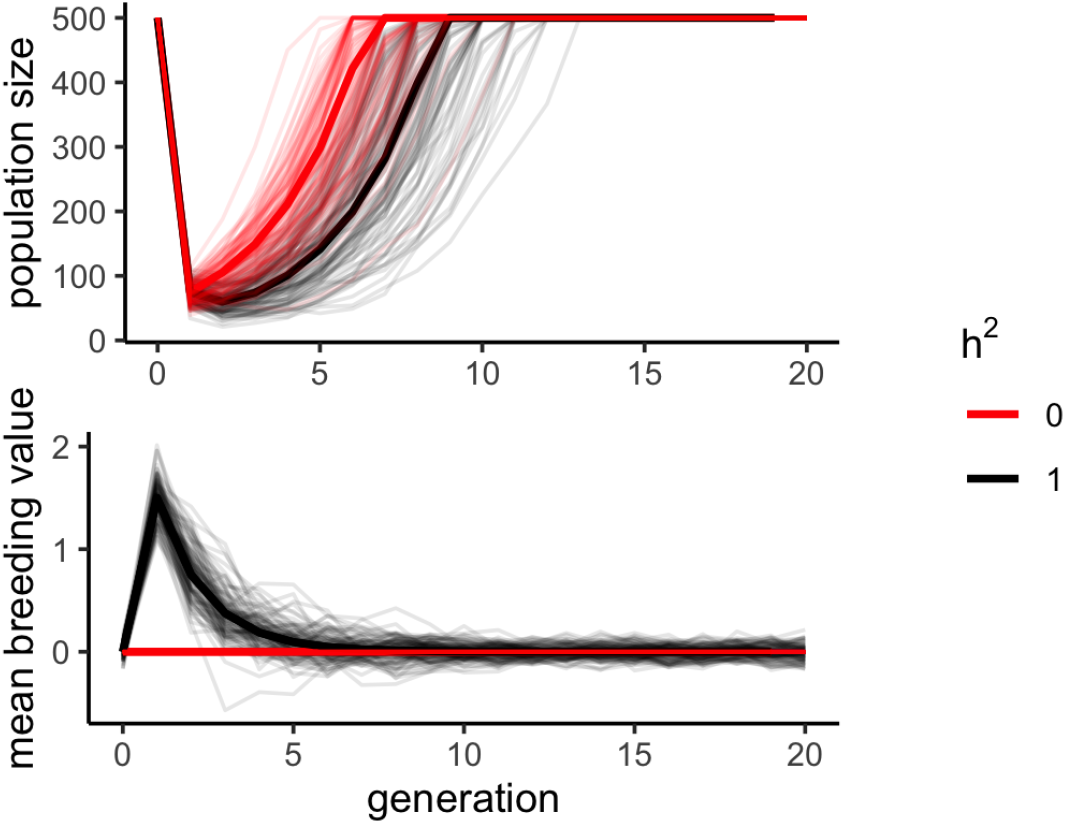
Population size over time for populations with *h*^2^ = 0 (red) and *h*^2^ = 1 (black) after a single-generation extreme event of size Δ*θ* = 3. Phenotypic variance is the same for both populations 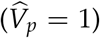. Faded lines are 100 simulations and solid lines are the model predictions using Equations (2) and (4). Parameters: *ω* = 1, *λ* = 2. Red: *V*_0_ = 0, *V_e_* = 1, Black: *V*_0_ = 3/4, *V_e_* = 0.

Previous work on evolution in changing environments can provide intuition for how evolution might affect extinction risk during or after a pulse disturbance. One focus of the evolutionary rescue literature has been on understanding the consequences of phenotypic change in the context of a sudden, long-term or permanent environmental shift (a press perturbation). These studies, some of which account for demographic stochasticity, underline the importance of genetic variance for increasing the probability of rescue (Gomulkiewicz and Holt, 1995, reviewed in Alexander et al., 2014; Bell, 2017). That is, populations that are able to adapt rapidly to the new environment have a higher chance of persisting. Similarly, studies of adaptation in fluctuating environments suggest that if an environment is predictable, such as the case of positively autocorrelated fluctuations, genetic variation reduces lag load (Charlesworth, 1993; Lande and Shannon, 1996; Chevin, 2013). This reduction in the lag load leads to higher population percapita growth rates and, consequently, is expected to reduce extinction risk. Whereas, when the environment is unpredictable, genetic variance typically increases the lag load. These studies calculated lag load and per-capita growth rates in a number of environmental contexts including a press perturbation, randomly fluctuating environments, and cyclic environments. However, they did not account for demographic stochasticity, the ultimate cause of extinction. Furthermore, we explore the impact of phenotypic variance on the probability of persistence, which has not been emphasized as much in previous studies.

To understand extinction risk during and following a pulse disturbance, we introduce an individual-based model that fuses population demography with quantitative genetics. Using a mixture of computational and analytical methods, we examine how phenotypic variation and the heritability of this variation influences population growth, lag load, and extinction risk during and following a pulse perturbation. Moreover, we examine how the magnitude and direction of these effects depend on the duration and intensity of the pulse perturbation.

## Model

We use an individual-based model that combines the infinitesimal-model of an evolving quantitative trait with density-dependent demography. To gain insights beyond simulating the model, we derive analytical approximations of the probability of extinction using a mixture of deterministic recursion equations and branching process theory (Harris, 1964). We assume discrete, non-overlapping generations. The life cycle starts with viability selection. In each generation *t*, we impose stabilizing selection around some optimal trait value *θ_t_*, which is set by the environment in that generation, by making the probability of survival

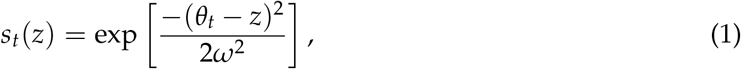

a Gaussian function of phenotype, z, with a strength of selection proportional to 1/*ω*^2^.

Following viability selection, survivors are randomly drawn with replacement to form mating pairs. Each pair then produces a Poisson number of offspring with mean 2*λ*. The population lives in a habitat that supports at most *K* individuals. Hence, if more than *K* offspring are produced, *K* are randomly chosen without replacement. The genetics of the population follows the infinitesimal model in which breeding values are determined by many loci of small effect (Fisher, 1918; Turelli, 2017). Under this model, an offspring’s breeding value is a draw from a normal distribution centered on the mean of its parents’ breeding values and with segregation variance *V*_0_ (which we assume is a constant). Its phenotype, *z*, is this breeding value, *g*, plus a random environmental component, *e*, which is a draw from a normal distribution with mean 0 and variance *V_e_*. We ignore dominance and epistasis, thus the phenotypic variance in generation *t* is the additive genetic variance plus the environmental variance, *V_p,t_* = *V_g,t_* + *V_e_*. At equilibrium, 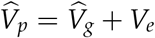.

Prior to experiencing an extreme event, the populations in the individual-based simulations start with a 100-generation burn-in from an initial state where all *N* = *K* individuals have breeding value *θ* = 0 and the optimal trait value *θ_t_* equals 0 throughout this period. The 100 generation burn-in is sufficiently long to ensure the model reaches a quasi-stationary state (Supplementary Figure S1). To model the extreme event of length *τ* after the burn-in period from generation −100 to 0, we increase the optimum trait value by Δ*θ* and revert it back to its original value after *τ* generations. For example, in a single-generation event, the optimum trait value changes before selection in generation 1 and then reverts back before selection in generation 2 (Figure 1). Unless otherwise stated, we use the parameter values *ω^2^* = 1, 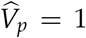, Δ*θ* = 3, *λ* = 2, and *K* = 500. These *ω^2^* and 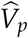 values represent strong selection and large phenotypic variance relative to those estimated in Turelli (1984), which we use to show the qualitative effect of variance load. Reducing the strength of selection (or decreasing the phenotypic variance) does not otherwise change our qualitative results. The values for Δ*θ* scale the strength of selection. For this set of parameter values, the optimum shift (Δ*θ* = 3) corresponds to three standard deviations beyond the mean of the trait distribution, and consequently, roughly 99.5 percent of the trait distribution will initially be smaller than the optimum and we expect roughly 80% of the population to die in the first generation. We have chosen a high growth rate, *λ*, to reduce extinction from demographic stochasticity in the absence of disturbance. We have chosen a large enough starting population size and carrying capacity, *K* = 500, to make approximations reasonable (e.g. normal distribution of traits).

## Approximations

### Approximating the evolutionary and population size dynamics

In Appendix A (see the supplementary Mathematica file for more details), we derive deterministic approximations for the dynamics of the mean breeding value 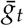, genetic variance *V_g,t_*, and population size *N_t_*. Briefly, if we assume the distribution of breeding values remains normally distributed, then we know the whole phenotypic distribution by tracking the mean and variance in the breeding values. Given the mean and variance in a given generation, we can then calculate the mean and variance in the next generation,

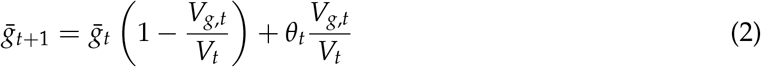

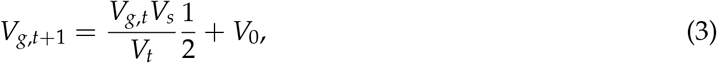

where *V_t_* = *V_g,t_* + *V_s_*, *V_s_* = *ω*^2^ + *V_e_* is the inverse of the effective strength of selection, and *V*_0_ is the variance in breeding values among siblings. We can also calculate the population size in the next generation

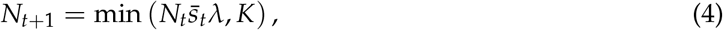

where the mean survival probability, 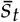, is calculated by integrating Equation (1) over the distribution of phenotypes in the population,

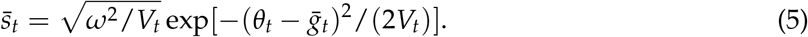

Regardless of the trait or environmental dynamics, the genetic variance approaches an equilibrium 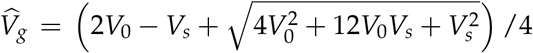 which increases with segregation variance and decreases with the strength of selection. In a constant environment, *θ_t_* = *θ* for all *t*, the mean breeding value approaches the optimum, 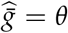, and, provided *λ* > 1, 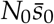 is large enough, and 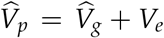 is small enough, the population size reaches carrying capacity, 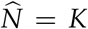. Starting from this equilibrium, we can then approximate the response of the population to a shift in the optimum using Equations (2)–(4).

### Approximating Extinction Risk

We next approximate the probability of extinction using branching processes (Harris, 1964). The probability generating function for the number of offspring produced by an individual with survival probability s is

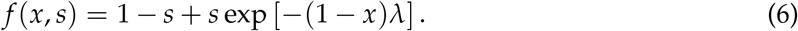

The probability of no offspring is *f* (0, *s*). Further, if *s*_1_,…, *S_N_t__* are the survival probabilities of the *N_t_* individuals in generation *t*, then the probability of extinction in generation *t* is 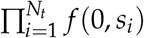. Here we approximate this by assuming all individuals in generation *t* have the average probability of survival, 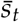, which is a reasonable approximation when the strength of selection is weak relative to the phenotypic variance. Defining 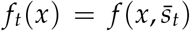, the probability of extinction in generation *t* is then simply *f_t_*(0)^*N_t_*^. Assuming that the effects of density-dependence are negligible from generation *t* to generation *T > t*, we can approximate the probability of extinction by the end of generation *T* as 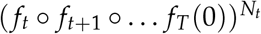, where ○ denotes function composition (Harris, 1964).

We take *t* = 1 to be the first generation of the extreme event and assume the population begins at carrying capacity. For an extreme event of duration *τ*, we define

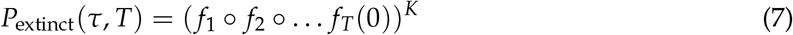

as our approximation for the probability of extinction by generation *T* since an extreme event of length *τ* began. To calculate the *s_t_* in Equation (7) we assume 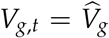 and use Equation (2) to get 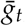, which together give 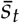 (Equation (1)).

## Results

### Demographic recovery

We first explore extreme events lasting a single generation. To characterize the impact of phenotypic variance and heritability on population size, we compare the demographic response of populations with low or high phenotypic variance, 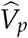, across a range of heritabilities, 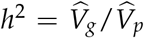. During the event, heritability has no effect on population size (compare black and red curves in 2A). In contrast, we see that phenotypic variation can have a large effect. A population with high phenotypic variance (thick gray curve) has a smaller population size than one with low phenoytpic variance (thin gray curve) immediately following a low severity extreme event, but a higher population size following more severe events. We also see this effect in the generation after the event (Figure 2B,C). This pattern stems from the dual role of phenotypic variance, in that it both increases variance load and contributes individuals with extreme traits who are then able to survive an extreme event. High phenotypic variance therefore reduces both mean fitness within a generation and the variance in fitness across generations – a form of short-term bethedging which can increase the geometric mean of fitness in the generations during and after the the disturbance event. The positive effect of bet hedging is seen when the event is severe and variation means more individuals on the tail of the distribution will survive the event.

**Figure 2:**
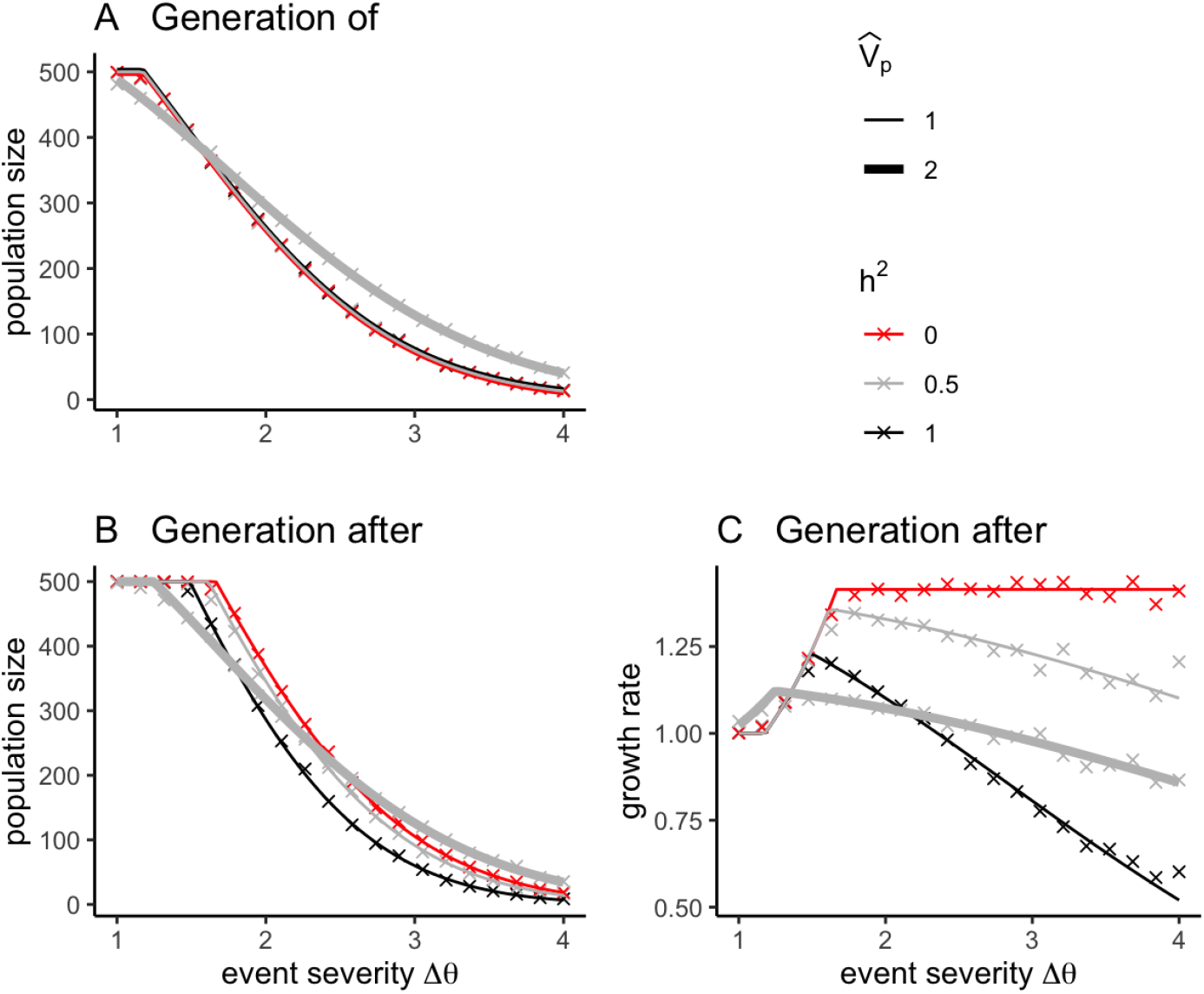
Population size response during the generation of a single-generation extreme event (A), the generation after the event (B), and the growth rate 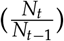 calculated as the population size in the generation after the event divided by the population size in the generation of (C) shown over a range of event severities Δ*θ*. Expectations using Equation (4) as curves and simulation results (mean of 100 replicates) as crosses. Parameters: *ω* = 1, *λ* = 2. Red: *V*_0_ = 0, *V_e_* = 1. Thick gray: *V*_0_ = 2/3, *V_e_* = 1. Thin gray: *V*_0_ = 5/16, *V_e_* = 1/2. Black: *V*_0_ = 3/4, *V_e_* = 0.

While heritability has no effect on survival during the event, it has a strong effect on population recovery in subsequent generations. In particular, heritability dampens the growth rate 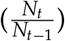 in subsequent generations (Figure 2C) as evolution in the generation of the event induces future maladaptation. This explains why in the generation after the event increasing segregation variance increases population size (thick gray curve crosses red near Δ*θ* = 3 in panel B) at a higher severity than the point which increasing environmental variance becomes beneficial (thick gray curve crosses black near Δ*θ* = 2 in panel B). The maladaptation induced by heritability continues past the generation after the event, generally slowing population recovery (Figure 1). In conclusion, phenotypic variance can be beneficial for population growth under single-generation severe events, but heritability is generally deleterious.

### Extinction Risk

When a single-generation extreme event is severe enough, increasing phenotypic variation lowers extinction risk both during and after the event (compare thick and thin gray curves in Figure 3A,C). The biological intuition behind this pattern is the same as in Figure 2A, where increased variance means more individuals survive the extreme event. However, at such large population sizes the extinction risk is essentially zero during a mild event. In other words, while having too much variance leads to considerable reduction in population size when events are mild, it is very unlikely to lead to extinction unless there is extremely high phenotypic variance or if carrying capacity is very low. In the former case load will cause extinction in the absence of extreme events (Supplementary Figure S2).

**Figure 3:**
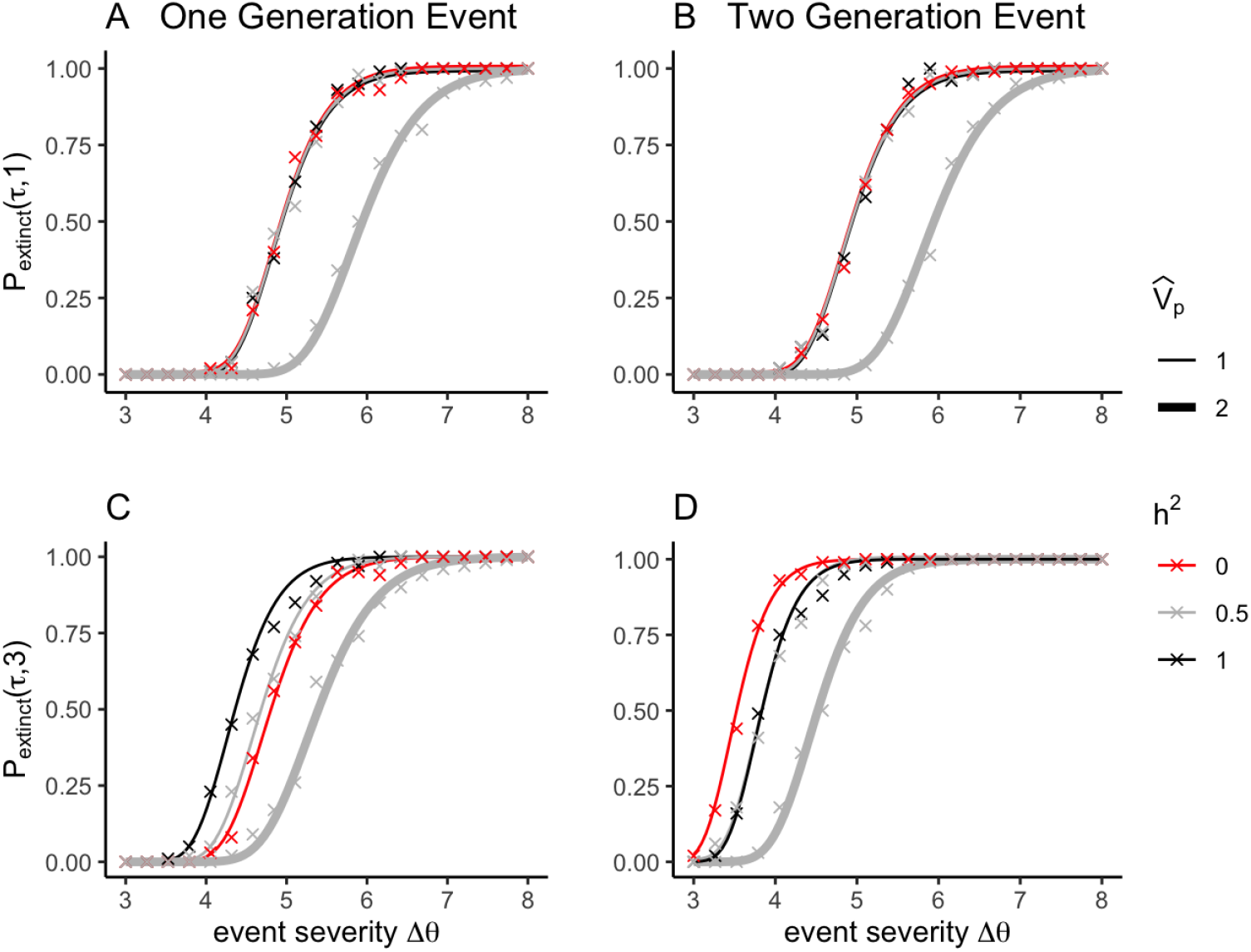
Extinction risk across increasingly severe events in the first generation of an extreme event, (A,B) and two generations later, (C,D). In A and C, the extreme event persists for a single generation, and in B and D, the extreme event persists for two generations. Expectations using Equation (7) as curves and simulation results (mean of 100 replicates) as crosses. Parameters: *ω* = 1, *λ* = 2. Red: *V*_0_ = 0, *V_e_* = 1. Thick gray: *V*_0_ = 2/3, *V_e_* = 1. Thin gray: *V*_0_ = 5/16, *V_e_* = 1/2. Black: *V*_0_ = 3/4, *V_e_* = 0.

Next, we compare populations with the same phenotypic variance but different heritabilities, to control for the effect of variance (i.e., variance load and bet hedging) and isolate the effect of evolution (compare black and red in Figure 3). When the extreme event lasts only one generation (Figure 3A,C), hertiability increases extinction risk in the generation following moderately severe extreme events. There is little effect when events are very mild or incredibly strong. Whereas, when an extreme event lasts two generations (Figure 3B,D), heritability reduces the risk of extinction in the generation following a moderately severe extreme event.

Finally, we explored how extinction risk varies across time for one-to four-generation moderately (Δ*θ* = 3.5) extreme events across a range of heritabilities. For single generation events, long-term extinction risk (10,000 generations) increases with heritability (Figure 4A), for the reasons above. However, for two generation events, long-term extinction risk is lowest at intermediate heritabilities (Figure 4B). And for three and four generation events, long-term extinction risk decreases with heritability (Figure 4C,D). These patterns hold for milder (Δ*θ* = 2.5) and more severe (Δ*θ* = 4.5) extreme events (Supplementary Figures S3–S4).

**Figure 4:**
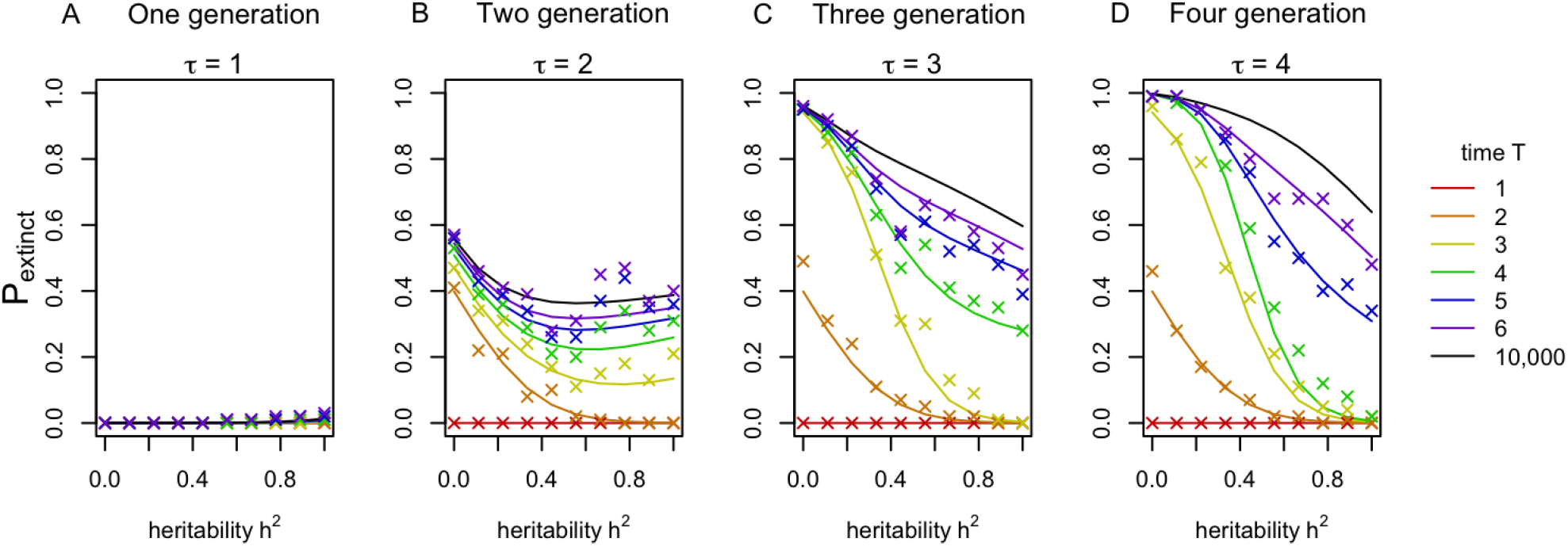
Extinction risk through time T across a range of heritability for extreme events lasting 1, 2, 3, or 4 generations. Time starts the generation the event began. Parameters: 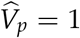, *ω* = 1, *λ* = 2, Δ*θ* = 3.5. Expectations using Equation (7) as curves and simulation results (mean of 100 replicates) as crosses.

While Equation (7) gives a good approximation of extinction risk, the function itself is too complex to give us intuition. Next, by writing down the geometric mean fitness of a population, we reproduce the general trends in long-term extinction risk, but with added clarity for how maladaptation contributes to these outcomes.

### Contribution of Lag Load

To better understand how evolution affects the probability of extinction, we approximate the geometric mean fitness of a population under the assumption that the genetic variance remains at the equilibrium value, 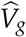, as expected based on Equation (3). If the extreme event lasts *τ* generations, then the geometric mean of fitness after *T* > *τ* generations is

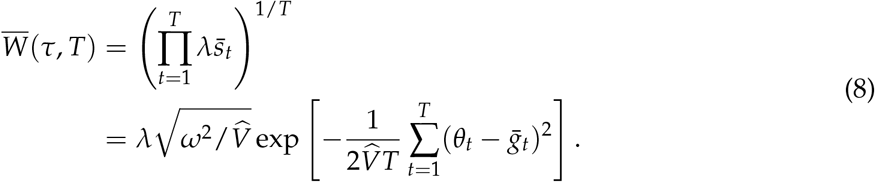

From Equation (8) we see that geometric mean fitness depends on the cumulative lag load, 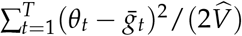.

Using Equation (2), we show in Appendix C that the cumulative lag load over *T* > *τ* generations for an event of length *τ* is

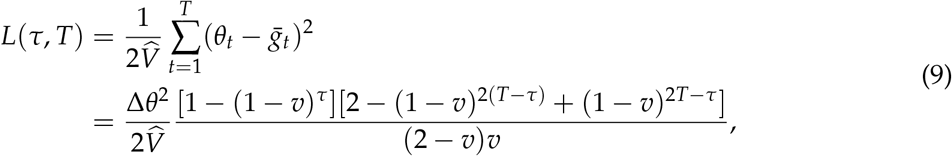

where 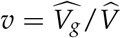 is a measure of evolvability (see Equation (2) and, e.g., equation 1 in Charlesworth 1993).

Taking the limit as time, *T*, goes to infinity, the cumulative lag load is

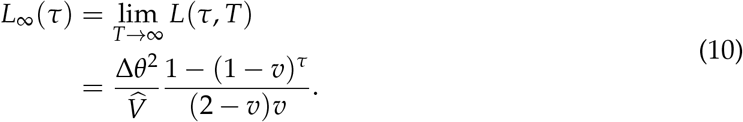

Generally, increasing the event length or increasing the event severity increases the cumulative lag load.

As 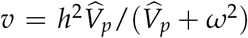, we can use Equation (10) to determine how heritability affects the cumulative lag load in the long term (Figure 5), holding 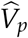 and *ω*^2^ (and thus the variance load) constant. When the extreme event only lasts one generation (*τ* = 1), the cumulative lag load equals 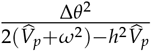. Hence, increasing heritability while holding 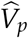 constant increases the cumulative lag load (solid purple curve in Figure 5), a trend consistent with the extinction probabilities for *τ* = 1 (Figure 4A). Alternatively, when the extreme event lasts two generations (*τ* = 2), the cumulative lag load equals 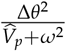 and is therefore independent of heritability when 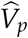 and *ω*^2^ are held constant (solid pink curve in Figure 5). Finally, when the extreme event lasts for more than two generations, the cumulative lag load is a decreasing function of heritability (yellow and green curves in Figure 5), a trend consistent with extinction probabilities decreasing with heritability when *τ* ≥ 3 (Figure 4C,D).

**Figure 5:**
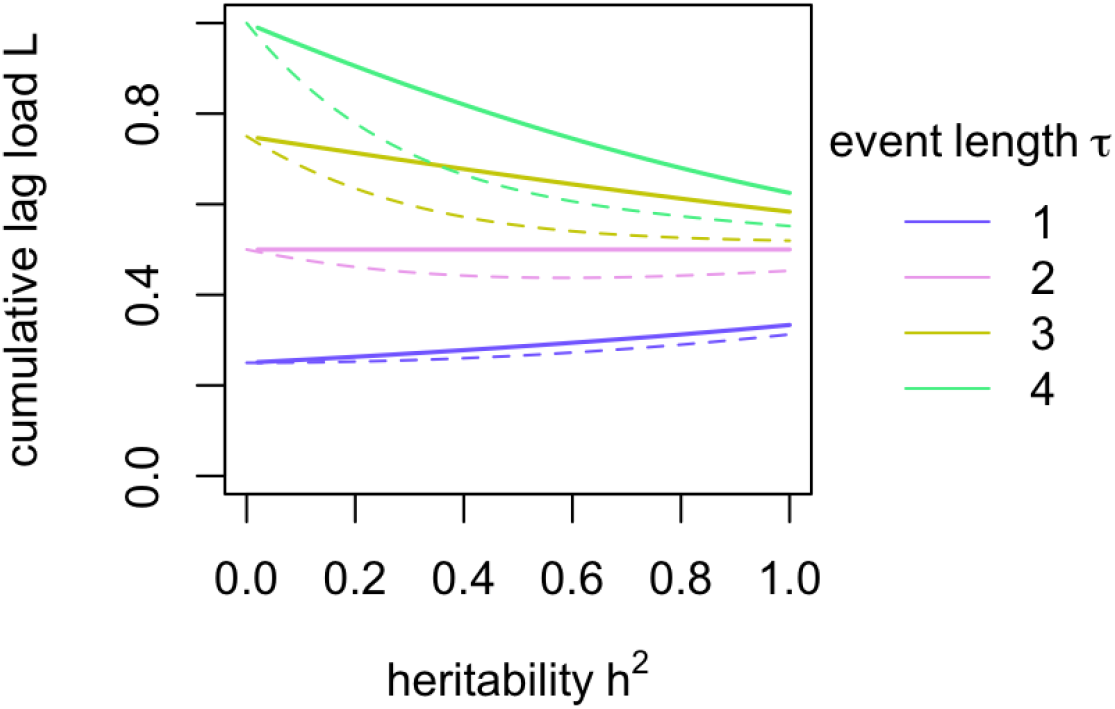
Cumulative lag load as a function of heritability. Dashed curves are Equation (9) with T = *τ* + 1 and solid are Equation (10). Colors correspond to the length of the extreme event. Parameters: 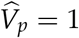, *ω* = 1, Δ*θ* = 1.

Taking the limit as both the time and event length go to infinity in Equation (9) and assuming weak selection (*V_s_* → ∞) we recover the cumulative lag load following a sudden non-reversing shift in the environment, 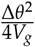 (equation 10 in Chevin, 2013). This is roughly half of Equation (10), since the environment never reverts back. Equation (9) can therefore be seen as a generalization of Chevin’s result to arbitrary time and event length under arbitrarily strong selection (provided our approximations in Appendix A hold).

## Discussion

Although it has been long recognized that evolution may affect a population’s response to a changing environment, previous studies have primarily focused on understanding this effect over the long term in the context of a single non-reversing environmental shift (a press disturbance) or a constantly fluctuating environment. Here, we are concerned with the short-term effect (finite T) of a pulse disturbance on population growth and extinction risk. By allowing pulses to be of any duration, this allows us to connect our results with those of both the press disturbance (large *τ*) literature and the constantly changing environment literature while providing insights into the transient effects (small T) following a disturbance. Our results provide two general conclusions about the effect of trait variation and its heritability on population growth and extinction risk during and following a pulse disturbance. First, trait variance, whether it is heritable or not, is a double-edged sword: adding a variance load due to stabilizing selection, yet providing individuals with more extreme traits who can survive large shifts in the environment. Second, while variance can be useful in the generation of a severe event, if heritable it slows demographic recovery and can therefore increase extinction risk in the generations after the event.

### Phenotypic Variance

Phenotypic variance, whether heritable or not, can be beneficial or deleterious. A simultaneous reduction in the mean and variance in fitness during the generations immediately prior, during, and immediately after an extreme event can increase the geometric mean of fitness during this time frame (Figure 2). This increase occurs when disturbances are sufficiently severe, in which case phenotypic variation can serve as a kind of short-term bet-hedging strategy. In addition to its effect on the geometric mean across multiple generations, variation in survival rates due to phenotypic variation, in and of itself, reduces variation in the total number of offspring produced by the population (Kendall and Fox, 2002) and, thereby, lowers extinction risk (Lloyd-Smith et al., 2005). This effect contributes to phenotypic variation reducing extinction risk.

Prior studies of evolutionary rescue have emphasized the beneficial aspect of genetic variance, but not non-heritable phenotypic variance, in rescuing a population from an abrupt shift in environment. For example, for constantly fluctuating environments, Charlesworth (1993) found that higher genetic variance reduces lag load when environmental fluctuations are large and, thereby, increases the long-term geometric mean of fitness. Similarly, studies of a sudden or gradual directional environmental shift found that high genetic variance at the time of the environmental shift promotes rescue (Alexander et al., 2014; Barfield and Holt, 2016; Bell and Collins, 2008; Gomulkiewicz and Holt, 1995).Here, by teasing out the effects of heritability and phenotypic variance, we emphasize the costs and benefits of each.

### Heritability

Contrary to evolutionary rescue for populations experiencing a press-perturbation (Barfield and Holt, 2016; Gomulkiewicz and Holt, 1995), we find that heritability increases extinction risk when pulse perturbations only last a single generation. We can gain some intuition for why this is by considering the limiting cases of traits not evolving versus tracking the optimal trait perfectly with a one generation lag. When the population is adapted to the original environment, but does not evolve in response to the extreme event, it experiences a reduction in fitness for the duration *τ* of the extreme event. In contrast, when selection tracks the optimal trait with a one generation lag, the population experiences a reduction in fitness only in the first and last generation of the extreme event. Hence, when the extreme event lasts one generation, extinction risk is higher for the evolving populations and when the extreme event lasts more than two generations, extinction risk is higher for the non-evolving populations. A similar understanding can be gained by adapting a classic population genetic model of allele frequency change with time-varying selection (Dempster, 1955; Felsenstein, 1976, see Appendix D).

In general, the trends in short-term extinction risk are parallel to the lag load predictions (Figures 4 and 5). However, they differ in two ways. When the extreme event lasts exactly two generations, the non-evolving population experiences the reduction in fitness in successive generations while the evolving population experiences this reduction in alternate generations. Hence, the evolving population is slightly less likely to go extinct (see Appendix B). Also, when a population exhibits an intermediate amount of tracking of the optimum, the variance in survival from year to year is reduced and therefore can lower the overall extinction probability. This effect is especially apparent in the result of two year events (Figure 4B).

While previous studies continuously varying environments have focused on large populations in the long term, calculating lag load and growth rates when rare, they provide intuition for our results on short-term extinction risk after a one-time event. A single-generation extreme event functions most like a negatively autocorrelated fluctuating or randomly fluctuating environment, in that a strong genetic response to selection in one generation is likely maladaptive in the next generation (Benaïm and Schreiber, 2019; Charlesworth, 1993; Chevin, 2013; Lande and Shannon, 1996). However, an extreme event lasting three or more generations acts like a positively autocorrelated environment in that the environment is more predictable and hence evolvability is favored. Cyclic oscillations with a high amplitude and long period would also fall into this category.

### Future Challenges and Directions

Our models include a number of simplifications to both evolutionary and demographic processes. First, we do not model the erosion of genetic variance with decreasing population size, which is expected due to greater genetic drift in smaller populations (Barfield and Holt, 2016; Lande and Barrowclough, 1987). Furthermore, we have limited our analysis to truly quantitative genetic traits (i.e. infinitely many small-effect alleles) where adaptation is not mutation-limited and evolution is easily reversed. Different genetic architectures, such as a few loci of large effect, likely will respond differently (Barghi et al., 2020). Second, in our model, the phenotypic variation due to environmental variation is random, which ignores the potential for phenotypic plasticity. Phenotypic plasticity has been shown to have variable effects on evolution and extinction risk that depend on the nature of environmental change (Kopp and Matuszewski, 2014; Lande, 2015). Third, we are only tracking a single trait, whereas extreme events likely select on many correlated traits. As genetic covariance can change the outcome of selection, further work is needed to explore the effects of multiple correlated traits. Fourth, we used the simplest possible model for density-dependence, the ceiling model, as used in previous evolutionary rescue studies (e.g., Bürger and Lynch, 1995). For other models of compensating density-dependence, such as the Beverton-Holt model (Beverton and Holt, 1957), we expect similar results. However, over-compensatory density-dependence, as seen in the Ricker model (Ricker, 1954), can result in oscillatory population-dynamics for which the timing of the extreme event relative to the oscillations may play a subtle role.

Our results call for the need of more empirical studies assessing trait and fitness changes after an extreme event has ended. The many case studies of evolution in response to extreme events focus on the adaptive nature of species responses in the short-term (e.g. Campbell-Staton et al., 2017; Coleman et al., 2020). What these studies often fail to mention is that this evolution can be maladaptive in the longer term. When the environment returns to normal, populations with shifted trait means could be worse off. To explore this effect, future empirical studies could be extended to track changes in trait values and population size over several generations following extreme events. For example, lizards can be tracked for several generations following a hurricane (Donihue et al., 2018) or a cold snap (Campbell-Staton et al., 2017). We highlight the Darwin’s finch example as one such study to do this (Grant and Grant, 2002), where finch traits and selection gradients were found to fluctuate in response to extreme events that lasted less than a generation. Larger beak sizes were selected for immediately following a drought due to the change in the seed composition, but in later years these beak sizes were maladaptive. However, because mean survival is higher in an average wet year regardless of beak size, the effect of this maladaptation on population recovery is not readily apparent. Ideally, this recovery could be compared to one causing similar mortality but less evolution.

An important next step will be to understand evolution and extinction risk under repeated extreme events. Extreme events, or catastrophic events, can be characterized by causing abrupt, infrequent, and large reductions in biomass or population size. Hence, prior work on adaptation and persistence using autoregressive processes to model environmental fluctuations (e.g., Benaïm and Schreiber, 2019; Charlesworth, 1993; Chevin, 2013; Lande and Shannon, 1996), do not accurately capture the nature of extreme events such as those presented in the continuous-time ecological models of Mangel and Tier (1994). We hope future studies exploring the impact of disturbance regime on evolution and extinction risk will benefit from the detailed understanding, like that provided here, of an evolving population’s response to a single extreme event.

## Appendix A Dynamics of the breeding value distribution and population size

Let the trait value of an individual be the sum of a genetic component (breeding value) and an environmental component, *z = g + e*. Assume we start with a population, in generation *t*, that has a normal distribution of breeding values, *p_g_*(*g, t*), with mean 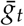 and variance *V_g,t_*. And assume each environmental component is independently chosen from a normal distribution, *p_e_*(*e*), with mean 0 and variance *V_e_*. The joint distribution of *g* and *e*, *p_ge_*(*g, e, t*), is then initially multivariate normal with mean 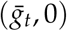, variances *V_g,t_* and *V_e_*, and no covariance.

Let the probability of survival for an individual with trait value *z* in generation *t* be

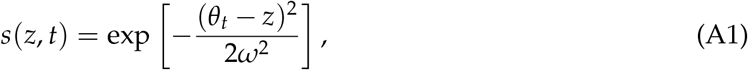

where *θ_t_* is the optimum trait value in generation *t* and 1/*ω*^2^ is the strength of selection. The joint distribution of *g* and *e* following viability selection is

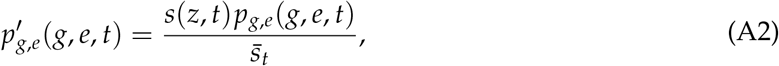

where

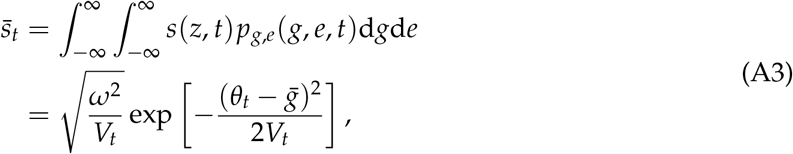

is the expected fraction of the population that survives in generation *t* (i.e., the population mean survival probability), with *V_t_ = V_g,t_ + V_s_* and *V_s_* = *ω*^2^ + *V_e_* the inverse of the effective strength of selection. Integrating over environmental effects then gives the distribution of breeding values amongst the survivors

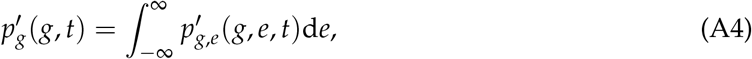

which is normal with mean 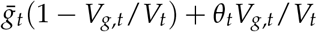 and variance *V_g,t_* (1 − *V_g,t_ /V_t_*). The mean breeding value is thus shifted towards *θ_t_* with a weight of *V_g,t_ /V_t_* and the genetic variance has been reduced by this fraction.

We next assume that the breeding value is determined by a large number of small effect loci, such that the distribution of breeding values amongst siblings, *p*_*g*,sibs_ (*g*|*g*_mid_), is normal with a mean equal to the midpoint of the parental breeding values, gmid, and a variance, *V*_0_, that does not depend on the parental genotypes or trait values (i.e., the infinitesimal model; Barton et al., 2017; Fisher, 1918). The distribution of breeding values among the offspring is then

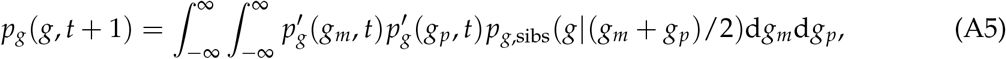

which is normal with mean

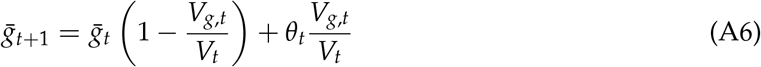

and variance

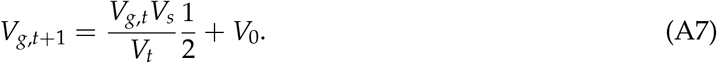

That is, the mean breeding value remains constant through reproduction while the variance before reproduction is first halved (due to essentially ”blending inheritance” between the parents) and then increased by segregation, *V*_0_.

So we see that given the initial distribution of breeding values is normal, with Gaussian selection the breeding value distribution remains normal, allowing us to track the entire distribution of breeding values (and therefore phenotypes) across generations by keeping track of only its mean and variance. The variance dynamics are independent of the environment (*θ_t_*) and the breeding values; solving Equation (A7) gives the genetic variance in generation any t. This expression is rather complicated (see Mathematica file), however it reaches an equilibrium

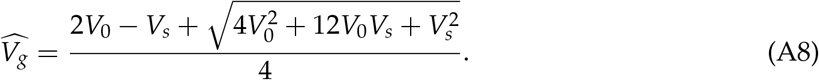

Holding genetic variance constant at its equilibrium (which is reasonable given the variance is not expected to change with the environment or breeding values), in a constant environment, *θ_t_* = *θ*, the mean breeding value in any generation t is found by solving Equation (A6),

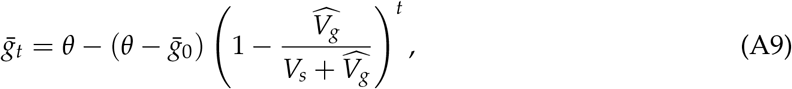

implying a geometric approach to 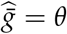 that becomes faster with 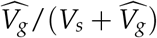.

We assume each individual that survives viability selection produces *λ* offspring, and that if more than *K* offspring are produced then *K* of these are randomly chosen to start the next generation. If the population size in generation *t* was *N_t_* then the population size in generation *t* + 1 is expected to be

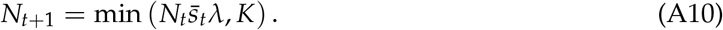

## Appendix B Extinction Risk in Single and Two Generation Events

In this Appendix, we examine the effect of long-term extinction risk when populations are either not evolving or are perfectly tracking, with a one-generation lag behind the optimal trait value. Let *s_o_* and *s_m_* be the survivorship of individuals with the optimal trait or the maladaptive trait. The offspring probability generating functions for these individuals are *f_o_*(*x*) = *f*(*x, s_o_*) and *f_m_*(*x*) = *f*(*x, s*_m_), respectively, where *f*(*x,s*) = 1 — *s* + *s*exp(*λ*(1 − *x*)). Let 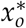 and 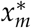 be the asymptotic extinction probability for the lineage of a single individual if it always exhibits the optimal trait and if it always is maladapted, respectively. Namely, 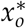 and 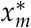 are the smallest fixed points of *f_o_* and *f_m_*, respectively, on the interval 0 ≤ *x* ≤ 1.

If a disturbance event lasts *τ* ≥ 1 generations, then the eventual extinction probability of the lineage of a non-evolving individual equals

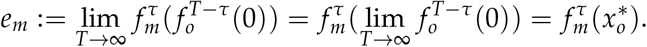

While the eventual extinction probability of the lineage of an individual with a one-generation lagged tracking of the optimal trait equals

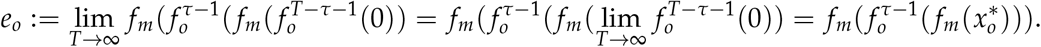

As *s_o_* > *s_m_*, we have *f_o_*(*x*) < *f_m_*(*x*) for all 0 ≤ *x* < 1, and 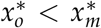. Furthermore, *f_i_*(*x*) are strictly increasing functions of *x*, *f_i_*(*x*) > *x* for 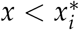, and *f_i_*(*x*) < *x* for 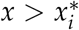 for *i = o, m*. Now suppose *τ* = 1. Then 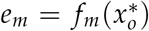 and 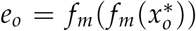. As 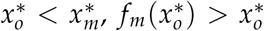. As *f_m_* is an increasing function, it follows that 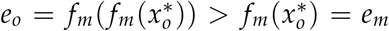. Now suppose *τ* = 2. Then 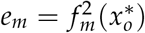 and 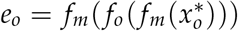. As 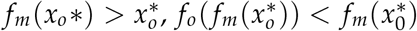. As *f_m_* is increasing, it follows that 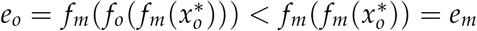.

## Appendix C Cumulative lag load

Here we show how to derive Equation (9). Our goal is to develop a formula for the cumulative squared displacement, 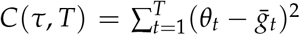, given event length *τ*. First note that Equation (2) implies that, with constant genetic variance 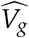, the mean trait displacement in the next generation is *g*_*t*+1_ − *θ*_*t*+1_ = (*g_t_* − *θ_t_*)(1 − *v*), where 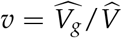 is a measure of evolvability. Thus, if the optimum is fixed at some arbitrary value for *τ* generations then the displacement in generation *t*, *d_t_* = *g_t_* − *θ_t_*, is *d_t_* = *d*_0_(1 − *v*)^t^ and the cumulative squared displacement over those *τ* generations is 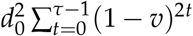. If the optimum then reverts to its original value for a further *T* − *τ >* 0 generations then the initial displacement is *d*_0_(1 − *v*)^*τ*^ − *d*_0_ and the cumulative squared displacement over this period is 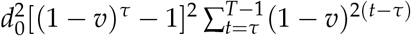. Combining these two sums we get

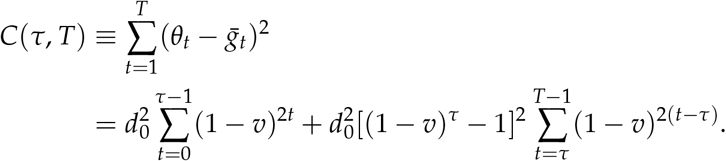

Multiplying by 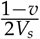, evaluating the sums, and setting the initial displacement as *d*_0_ = Δ*θ* gives Equation (9) in the main text.

## Appendix D Adapting a Population Genetic Model

To gain a better understanding of why cumulative lag load depends on event length, we adapt previous population genetic models of temporally variable selection. Consider a haploid case with the ratio of the initial frequencies of two alleles being 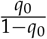. The ratio of the frequencies of the alleles at time *t* + 1, 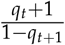, equals the product of the selection coefficients st from *t* = 0 to time *t* = *T*, multiplied by the ratio of the initial frequencies (Dempster, 1955; Felsenstein, 1976).

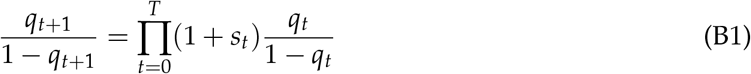

Here, rather than allele frequency change, we consider the product of fitnesses over one extreme event of length tau, where s is the selection coefficient corresponding to the change in the optimum during the event. When a population starting at the optimum perfectly tracks the extreme event, the product of fitnesses is

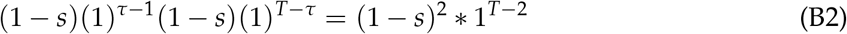

where fitness is reduced by s initially when the environment shifts to a new optimum, and then again when the environment returns to the original optimum. On the other hand, when a population starting at the optimum does not track the extreme event, the product of fitnesses is

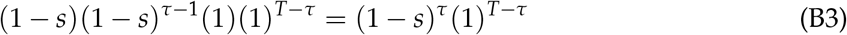

In the case of a two generation event, the product of fitnesses is (1 − *s*)^2^ regardless of whether it is a perfectly tracking population or a population that does not track the event. In events longer than two generations, perfectly tracking the environment is better.

## Supplementary Figures

**Figure S1:**
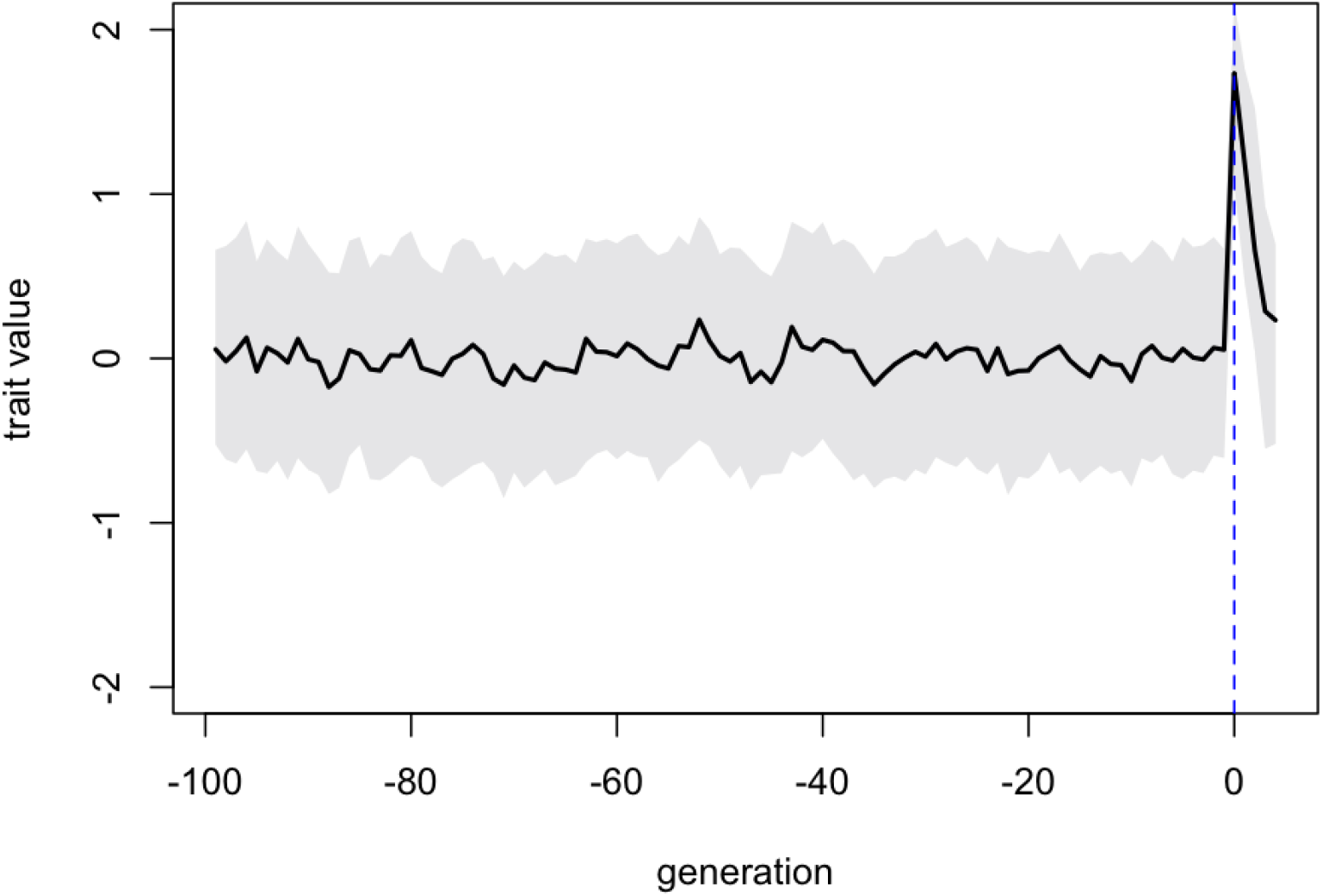
Rapid expansion and stabilization of phenotypic variance during the 100 generation burn-in with *V_e_* = 0, *V*_0_ = 1. Black line is mean trait value and gray shaded region extends from minimum to maximum trait values. The dashed blue curve indicates a one generation extreme event.

**Figure S2:**
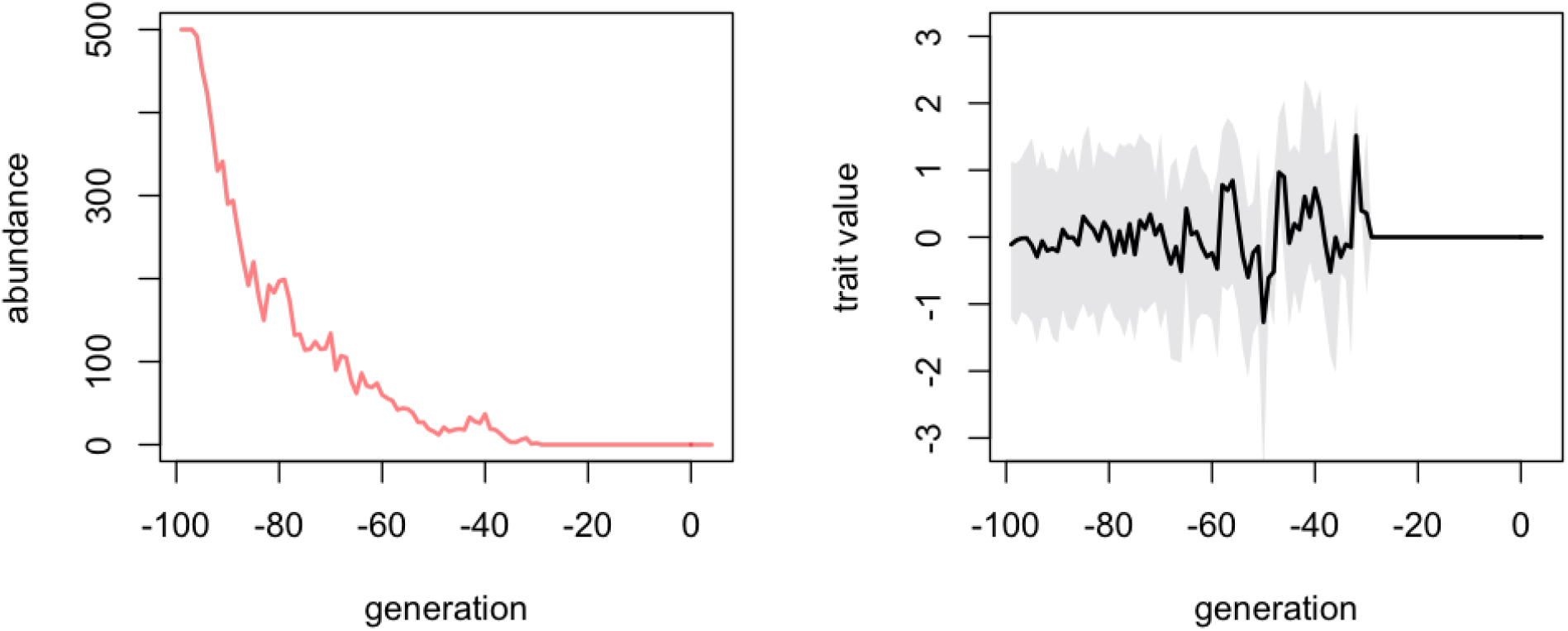
Extinction in a population with high variance load with *V*_0_ = 3, *V_e_* = 0. Black line is mean trait value, grey shaded region extends from minimum to maximum trait values.

**Figure S3:**
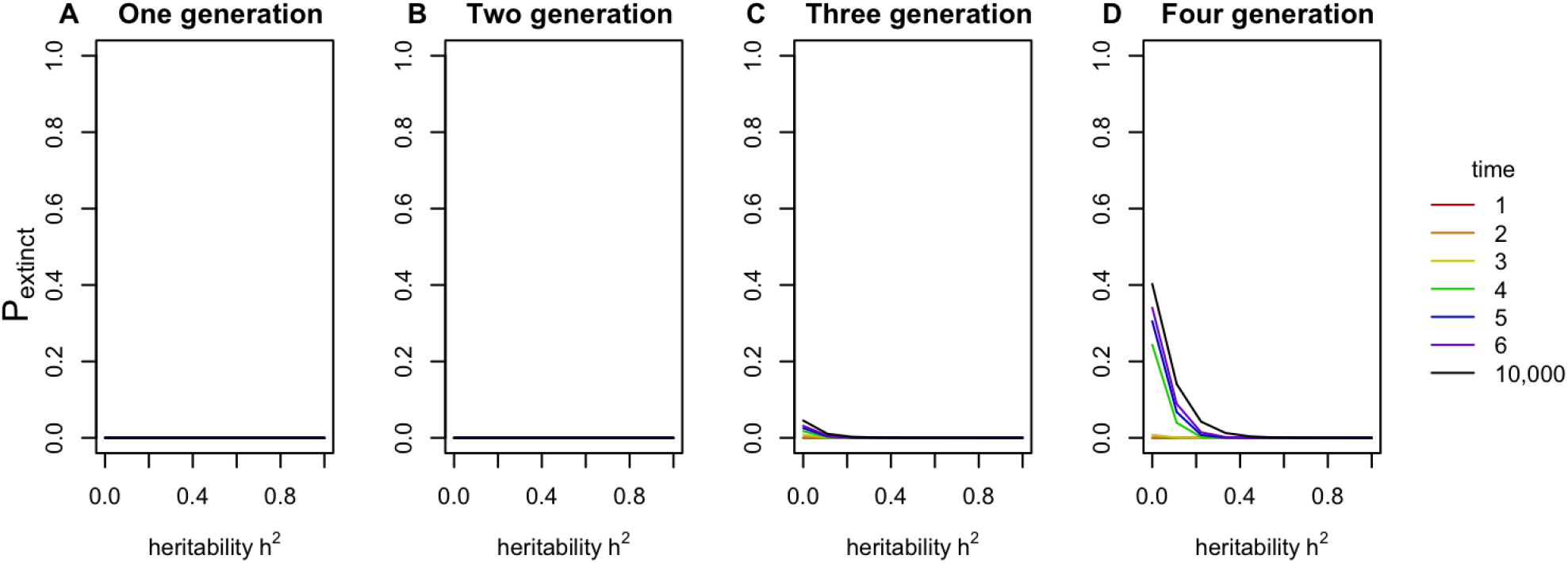
Extinction risk through time across a range of heritability for extreme events lasting 1, 2, 3, or 4 generations. Time starts the generation the event began. 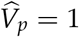, *ω* = 1, Δ*θ* = 2.5.

**Figure S4:**
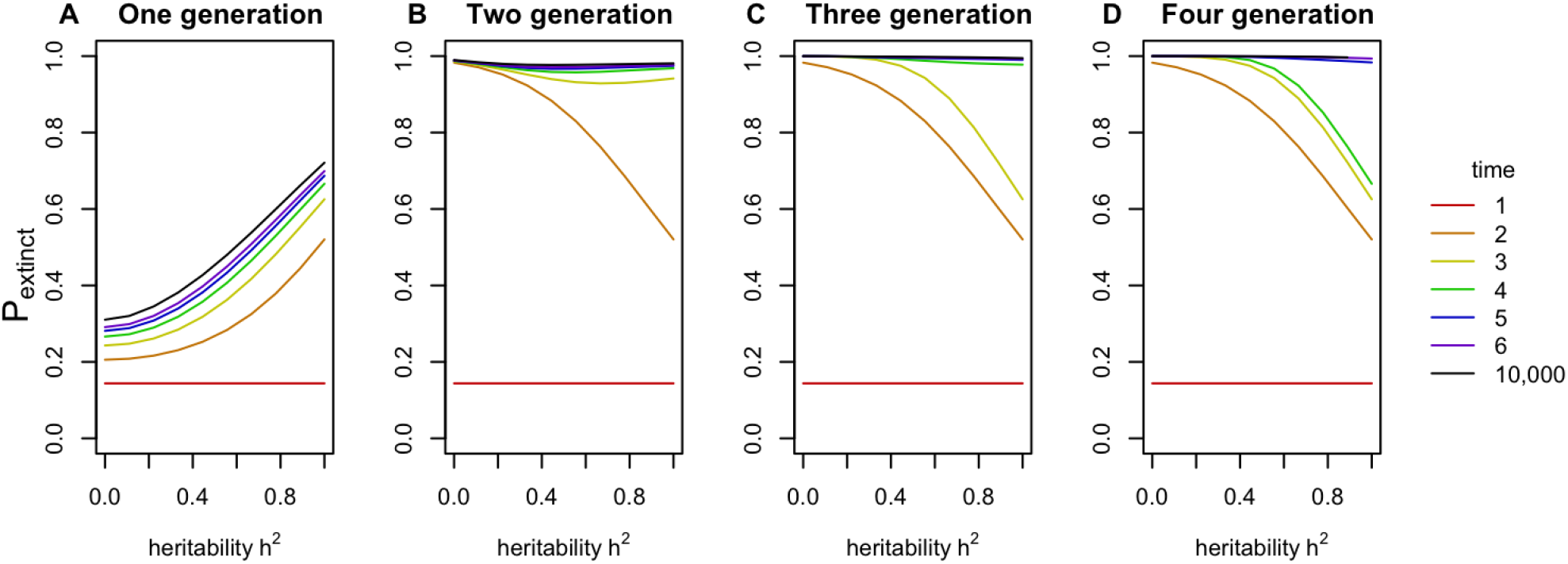
Extinction risk through time across a range of heritability for extreme events lasting 1, 2, 3, or 4 generations. Time starts the generation the event began. 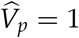, *V_s_* = 1, Δ*θ* = 4.5.

